# A group of nuclear factor Y transcription factors are likely sub-functionalized in the endosperm development of monocot

**DOI:** 10.1101/149047

**Authors:** E Zhiguo, Li Tingting, Zhang Huaya, Liu Zehou, Deng Hui, Sharma Sandeep, Wei Xuefeng, Wang Lei, Niu Baixiao, Chen Chen

## Abstract

Nuclear factor Y (NF-Y) is a heterotrimeric transcription factor that consists of three subunits, NF-YA, NF-YB, and NF-YC. Although NF-Ys play multiple roles in plant development, their functions during endosperm development are not well understood. In this study, we identified eight NF-Y encoding genes, including *OsNF-YA8, OsNF-YB1,9*, and *OsNF-YC8,9,10,11,12*, which predominantly express in the rice endosperm. Interestingly, the closest homologs of these OsNF-Ys are present only in the monocot species. All the genes are preferentially expressed in the endosperm, suggesting their roles in the regulation of endosperm development. A systemic analysis of the interactions between rice endosperm-preferential NF-Ys in yeast revealed that NF-YBs and NF-YCs could interact with each other. OsNF-YA8 is a recently evolved NF-YA in rice. Generally, NF-YA does not interact with NF-YB monomers in plants; however, in the present study, we found that OsNY-YA8 interacts with OsNF-YB9. Our results also indicated that the endosperm-preferential OsNF-YBs and OsNF-YCs could interact with some ethylene response factors (ERFs) of rice. Unlike the OsNF-YC8,9,10, the members of OsNF-YB1, 9 or OsNF-YC 11 and 12 showed lack of transcriptional activation when present alone. However, they displayed functional activity while in dimer form. In addition, *OsNF-YB1* knockout lines showed significant changes in the seed morphology, further confirms its role in endosperm development. Our findings have provided strong evidences that the group of phylogenetically conserved NF-Ys are differentiated in monocots to regulate the endosperm development.

## Introduction

Rice (*Oryza sativa*) provides most of the calories consumed by human beings globally. The endosperm is the edible part of the rice plant and other cereals. After double fertilization, the primary rice endosperm nucleus develops into the syncytium, a cell with multiple free nuclei that is formed by cell division but uncoupled with cytokinesis (Wu *et al.*, 2016). Simultaneous cellularization of the free nuclei initiates proliferation of the endosperm cells. After several rounds of mitotic division, the endosperm cells occupy the central vacuole and start to accumulate starch (Wu *et al.*, 2016). Two to five layers of the outermost cells of the endosperm are differentiated into an aleurone layer while the inner cells are differentiated into storage cells (Zhou *et al.*, 2013). The pattern of endosperm development in other monocots is similar to that of rice (Olsen, 2001; Leroux *et al.*, 2014; Zhang *et al.*, 2016*b*).

Several groups of transcription factors, such as the MADS-box genes *OsMADS6* (Zhang *et al.*, 2010), *OsMADS29* (Yin and Xue, 2012) and *OsMADS87* (Chen *et al.*, 2016), the bZIP family gene *RISBZ1* (Yamamoto *et al.*, 2006; Kawakatsu *et al.*, 2009), the DOF family gene *RPBF* (Yamamoto *et al.*, 2006; Kawakatsu *et al.*, 2009), and the WRKY family gene *OsWRKY78* (Zhang *et al.*, 2011), are essential for rice endosperm and seed development. The nuclear factor Y (NF-Y), also known as the Heme Activator Protein or CCAAT-binding factor, is conserved across kingdoms (Petroni *et al.*, 2012; Laloum *et al.*, 2013). The NF-Ys consist of three families: NF-YA, NF-YB, and NF-YC. Owing to possession of the Histone Fold Domains (HFDs), NF-YB and NF-YC can form a heterodimer, which generates a surface for NF-YA accession to form an NF-YA/B/C trimeric complex (Nardone *et al.*, 2016; Gnesutta *et al.*, 2017). The CCAAT-motif binding ability of NF-YA (Laloum *et al.*, 2013; Gnesutta *et al.*, 2017) and the transcriptional activation activities of NF-YB and NF-YC (Coustry *et al.*, 1995, 1996) allow the heterotrimeric complex to act as a transcription factor. This NF-Y complex may further interact with other transcription factors to regulate the expression of downstream targets (Yamamoto *et al.*, 2009; Cao *et al.*, 2014; Huang *et al.*, 2015; Xu *et al.*, 2016). Interestingly, there are only one or two encoding genes of each NF-Y family in mammals and yeast (Dolfini *et al.*, 2012). However, plants have substantially expanded their NF-Y genes (Laloum *et al.*, 2013). As an example, the rice genome encodes 11 NF-YAs, 11 NF-YBs, and 12 NF-YCs (Yang *et al.*, 2016). The expansion of plant NF-Ys may increase the number of possible NF-Y complexes and likely contributes to the neo-functionalization or sub-functionalization of the complexes. For instance, a group of phylogenetically related NF-Ys that predominantly expressed in nodules are specifically involved in nodulation in legume (Baudin *et al.*, 2015). Several NF-Ys have been found to be indispensable for seed development. *Leafy Cotyledon 1* (*LEC1*), also known as *NF-YB9* of *Arabidopsis* (*AtNF-YB9*), and its homolog *LEC1-like* (*L1L* or *AtNF-YB6*) are required for embryo maturation (Kwong *et al.*, 2003; Lee *et al.*, 2003). LEC1 and L1L belong to a phylogenically conserved clade of plants (Xie *et al.*, 2008). NF-YB7 of rice (OsNF-YB7) and OsNF-YB9 are the most similar homologs of LEC1 in rice. *OsNF-YB7/OsHAP3E* is expressed in the developing embryo and callus (Thirumurugan *et al.,* 2008). The ectopic expression of *OsNF-YB7* results in vegetative and reproductive development defects (Ito *et al.*, 2011; Zhang and Xue, 2013). However, due to the lethality caused by RNA-interference mediated gene silencing of *OsNF-YB7* (Ito *et al.*, 2011; Zhang and Xue, 2013), its role in embryogenesis and seed development remains to be elucidated. *OsNF-YB1*, an endosperm-specifically expressed gene, can coordinate with NF-YC members to regulate cell proliferation, grain filling, and sugar loading of the endosperm (Sun *et al.*, 2014; Bai *et al.*, 2015; Xu *et al.*, 2016). Bai et al. (2015) showed that OsNF-YB1 was able to recognize the CCAAT motifs of some sucrose transporters, while Xu et al. (2016) suggested that OsNF-YB1 likely lacks CCAAT-box binding activity but it could bind to ERF transcription factors in endosperms to regulate gene expression. In addition, OsNF-YB1 is able to interact with OsMADS18, a MADS-box family transcription factor of rice (Masiero *et al.*, 2002). The biological significance of the interaction is yet to be identified.

Some NF-Ys are found to be preferentially expressed in rice, wheat, and maize endosperms (Stephenson *et al.*, 2007; Yang *et al.*, 2016; Zhang *et al.*, 2016*a*). However, the importance of NF-Ys for endosperm development is not well understood. Here, we have performed comprehensive analysis with endosperm preferential NF-Ys in rice and identified two phylogenically-conserved NF-YB clades and one NF-YC clade in monocots. The endosperm specific gene expression and the interactions with other endosperm preferential NF-Y members suggest that these family genes have specific roles in endosperm development. The study provides novel insights into the functional role of NF-Ys in rice seed development.

## Results

### Endosperm-preferentially expressed NF-Ys in rice

The rice genome encodes 11 NF-YAs, 11 NF-YBs, and 12 NF-YCs (Yang *et al.*, 2016). A cluster analysis of the expression profiles showed that a group of NF-Ys, including *OsNF-YA8, OsNF-YB1*, and *OsNF-YB9*, and *OsNF-YC8* to *OsNF-YC12*, were predominantly expressed in rice endosperms (**Fig. 1 and Fig. 2A**). We further confirmed the expression pattern of these NF-Ys by real-time PCR. All endosperm-preferential NF-Ys were activated 3 days after fertilization (DAF) to 4 DAF (**Fig. 2B**). There is another rice NF-YB gene, *OsNF-YB7*, which also showed the seed preferentially-expressed pattern (**Fig. 1 and Fig. 2**). However, the microarray data (**Fig. 2A)** and a previous study (Thirumurugan *et al.*, 2008) suggested that its expression is mainly restricted to the embryo rather than the endosperm. The results show that each NF-Y family has members predominantly expressed in rice endosperms, which infers that the NF-Y complex may play important roles in the regulation of endosperm development.

**Figure 1.**
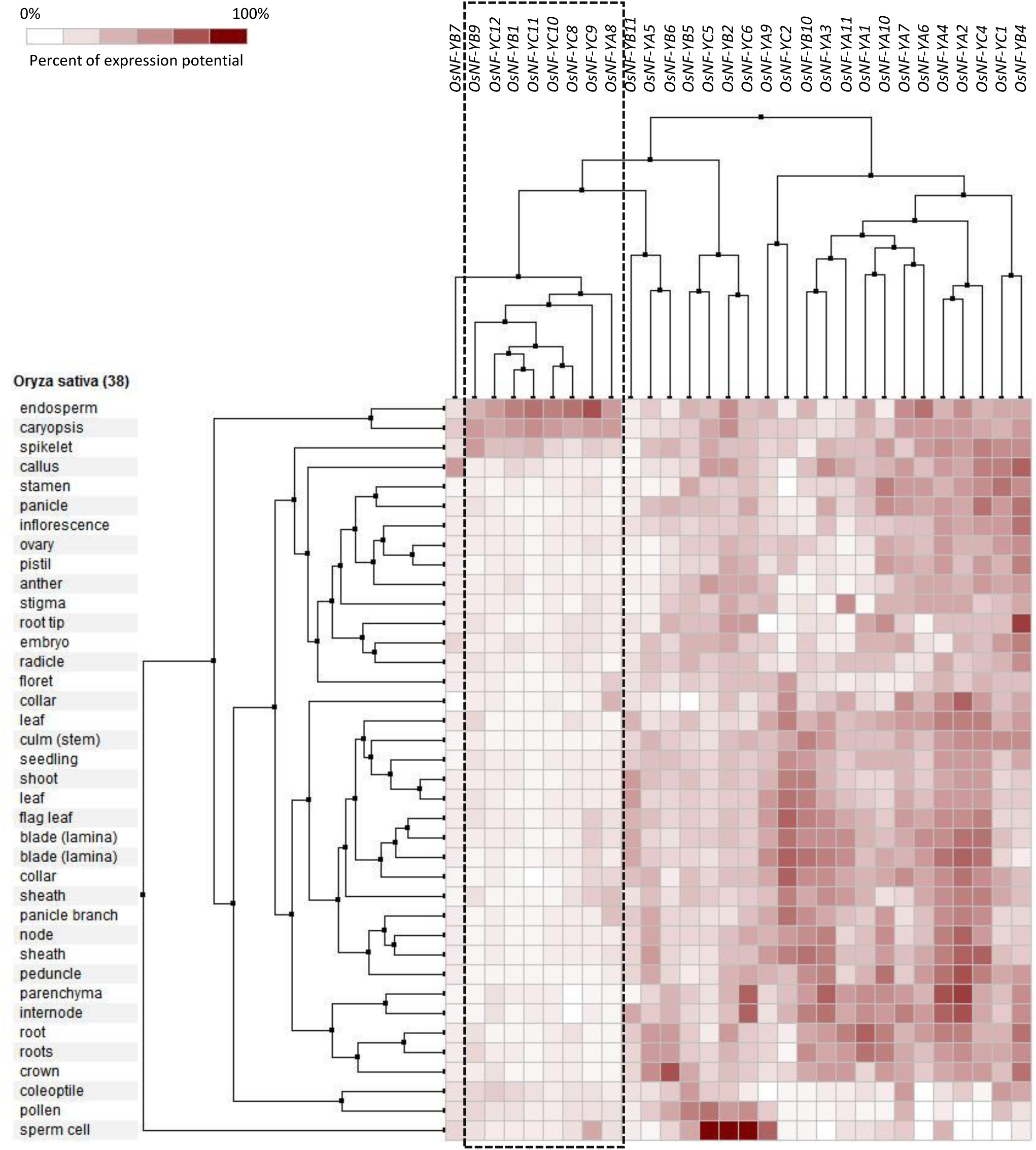
Cluster analysis of the expression of rice NF-Ys in various tissues. A group of rice NF-Ys (boxed by dash lines), including *OsNF-YA8, OsNF-YB1, OsNF-YB9, OsNF-YC8, OsNF-YC9, OsNF-YC10, OsNF-YC11* and *OsNF-YC12*, are predominantly expressed in the endosperm.

**Figure 2.**
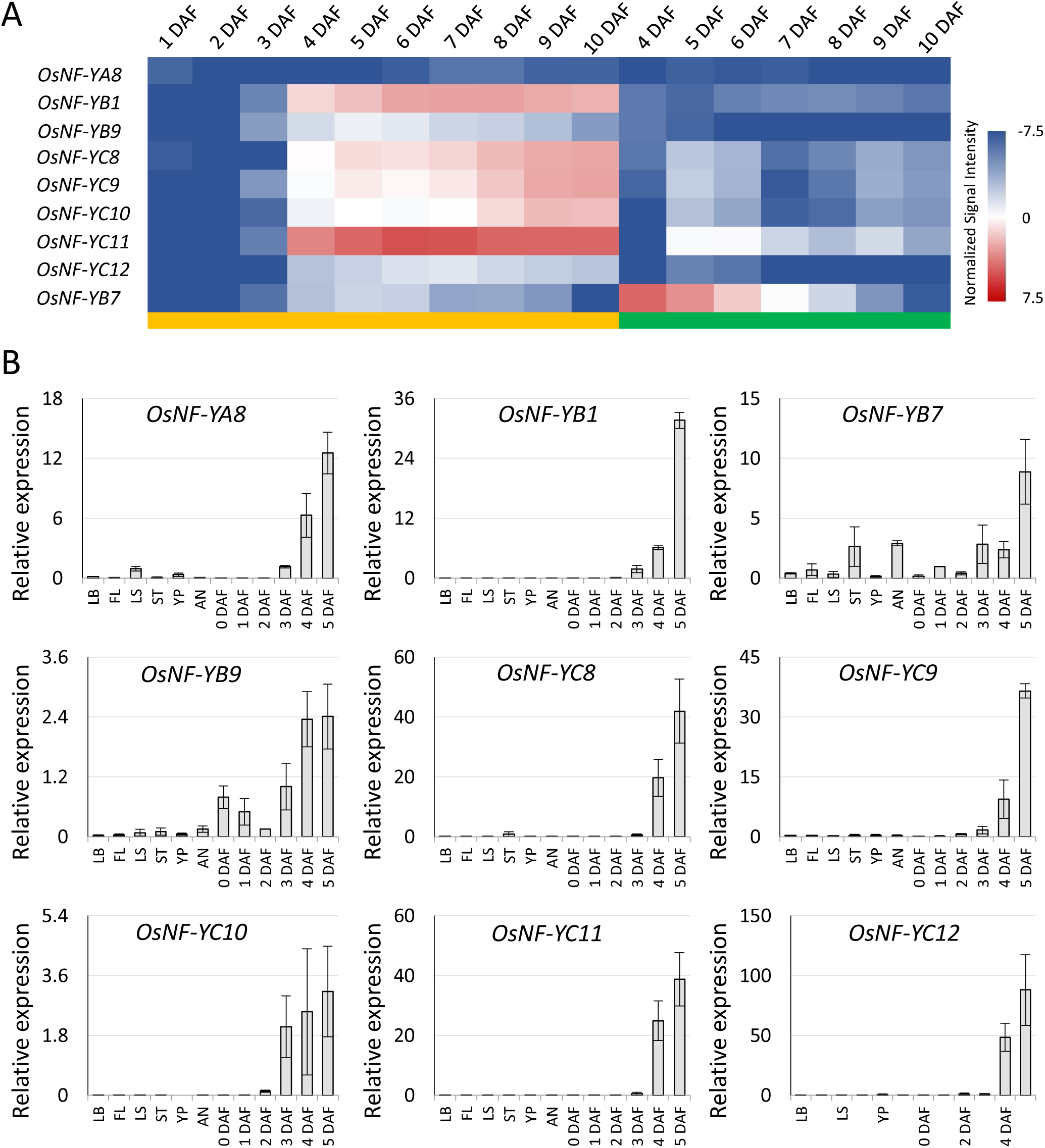
Activation of the seed-preferential *OsNF-Ys* after fertilization. **(A)** Heat map of the expression of *OsNF-YA8, OsNF-YB1, OsNF-YB7, OsNF-YB9, OsNF-YC8, OsNF-YC9, OsNF-YC10, OsNF-YC11* and *OsNF-YC12* at different days after fertilization (DAF). Yellow and green bars indicated ovary (including seed coat, endosperm and embryo) and embryo, respectively. **(B)** Confirmation of the seed-preferential expression pattern and gene activation after fertilization of *OsNF-YA8, OsNF-YB7, OsNF-YB1, OsNF-YB9, OsNF-YC8, OsNF-YC9, OsNF-YC10, OsNF-YC11* and *OsNF-YC12* by Real-time PCR. LB, FL, LS, ST, YP and AN indicate leaf blade, flag leaf, leaf sheath, stem, young panicle and anther, respectively; 0 DAF indicate unfertilized ovary; 1 to 5 DAF indicate ovaries of different ages (from 1 to 5 DAF). Three biological replicates are used for analysis; error bars indicate standard deviations.

### Phylogenetic analysis of endosperm-preferential NF-Y rice genes

To further study gene divergence of the endosperm-preferential NF-Ys, we performed a sequence alignment and phylogenetic analysis of all NF-YAs, NF-YBs, and NF-YCs in rice. Because the sequence outside the conserved domain is highly variable, we used the core sequence of the conserved domains for further analysis. In comparison with other NF-YAs, OsNF-YA8 showed several differences in the core sequence, many of which were located within the A2 domain **(Supplementary Figs. 1A).** It has been reported that the A2 domain is important for NF-YAs to bind with CCAAT motif (Nardone *et al.*, 2016). Due to the sequence variability in the A2 domain, it is possible that, in addition to the CCAAT motif, OsNF-YA8 may recognize other cis-elements. By using core sequence of OsNF-YA8 as a query search, we failed to identify its close homologs in other plant species which indicated that OsNF-YA8 might have evolved after the divergence of *Oryza* genus.

An alignment analysis of the NF-YB family members showed that the core sequence of OsNF-YB1 was distinct from the other NF-YBs of rice **(Supplementary Figs. 1B and E).** Many of these variations are presented within the α2 domain, which is believed to involved in NF-YA and NF-YC interactions with NF-YBs (Petroni *et al.*, 2012; Laloum *et al.*, 2013). The core sequences of OsNF-YB9 and OsNF-YB7, two LEC1 homologs in rice, are very similar **(Supplementary Fig. 1B),** although they expressed differently (**Fig. 2A**). A previous study has confirmed that the Asp at position 55 in LEC1 (or 84 in L1L) is diagnostic for LEC1 family members and is essential for gene function in *Arabidopsis* (Gnesutta *et al.*, 2017). Moreover, a crystal structural analysis suggested that His at position 79 was essential for the CCAAT-binding ability of L1L that compensate the substitutional changes of Lys→Asp at position 84 (Gnesutta *et al.*, 2017). The Asp and His residues are also conserved in OsNF-YB9 and OsNF-YB7 **(Supplementary Fig. 1B).** Since out of the core sequences are less conserved in OsNF-YB9 and OsNF-YB7, it keeps a question open whether the genes have similar functions in rice endosperm development. Interestingly, the LEC1 homologs in monocots can be divided into two phylogenetic clads. OsNF-YB7 is included in the same group of LEC1 and L1L, whereas OsNF-YB9 belongs to another clade that exclusively consists monocot homologs (**Fig. 3A**). Three substitutions were found by comparing the core sequences of the OsNF-YB7-like (OsYB7L) and OsNF-YB9-like (OsYB9L) proteins in different monocot species **(Supplementary Fig. 2).** The residue Thr at position 33 of OsYB7Ls was substituted by Ala, Val, or Leu, all are hydrophobic, in OsYB9Ls. The residues QREQ at position 49-52 of OsYB7Ls is much conserved, however, it is very variable in OsYB9Ls. In addition, the Tyr or Phe at position 89 of OsYB7Ls is substituted by Met in OsYB9Ls. Whether the variations in the genes sequence are responsible for their functional differentiations needs to be elucidated. In terms of the NF-YC family, OsNF-YC8, OsNF-YC9, OsNF-YC10, OsNF-YC11, and OsNF-YC12 are phylogenetically distinct from the other rice NF-YCs **(Supplementary Figs. 1C and F).** Even the intact nucleotide sequences of *OsNF-YC8, OsNF-YC9*, and *OsNF-YC10* are very similar **(Supplementary Fig. 3A),** indicating that these genes were formed by recent gene duplication events. Likewise, OsNF-YC11 and OsNF-Y12 show very high similarities **(Supplementary Fig. 3B).** Because of a very close relationship among OsNF-YC8, OsNF-YC9, and OsNF-YC10, and between OsNF-YC11 and OsNF-YC12, we used the core protein sequence of OsNF-YC8 and OsNF-YC12 to search their homologs in other plant species. The results showed that the homologs of OsNF-YC8 and OsNF-YC12 are only present in the monocots but not in the dicots (**Fig. 3B**), which clearly suggests that this group of genes are evolved after the divergence of dicots and monocots. Together, our results showed that the endosperm-preferentially expressed NF-YA, NF-YBs, or NF-YCs have very unique sequence features that separate them from widely-expressed canonical NF-Ys.

**Figure 3.**
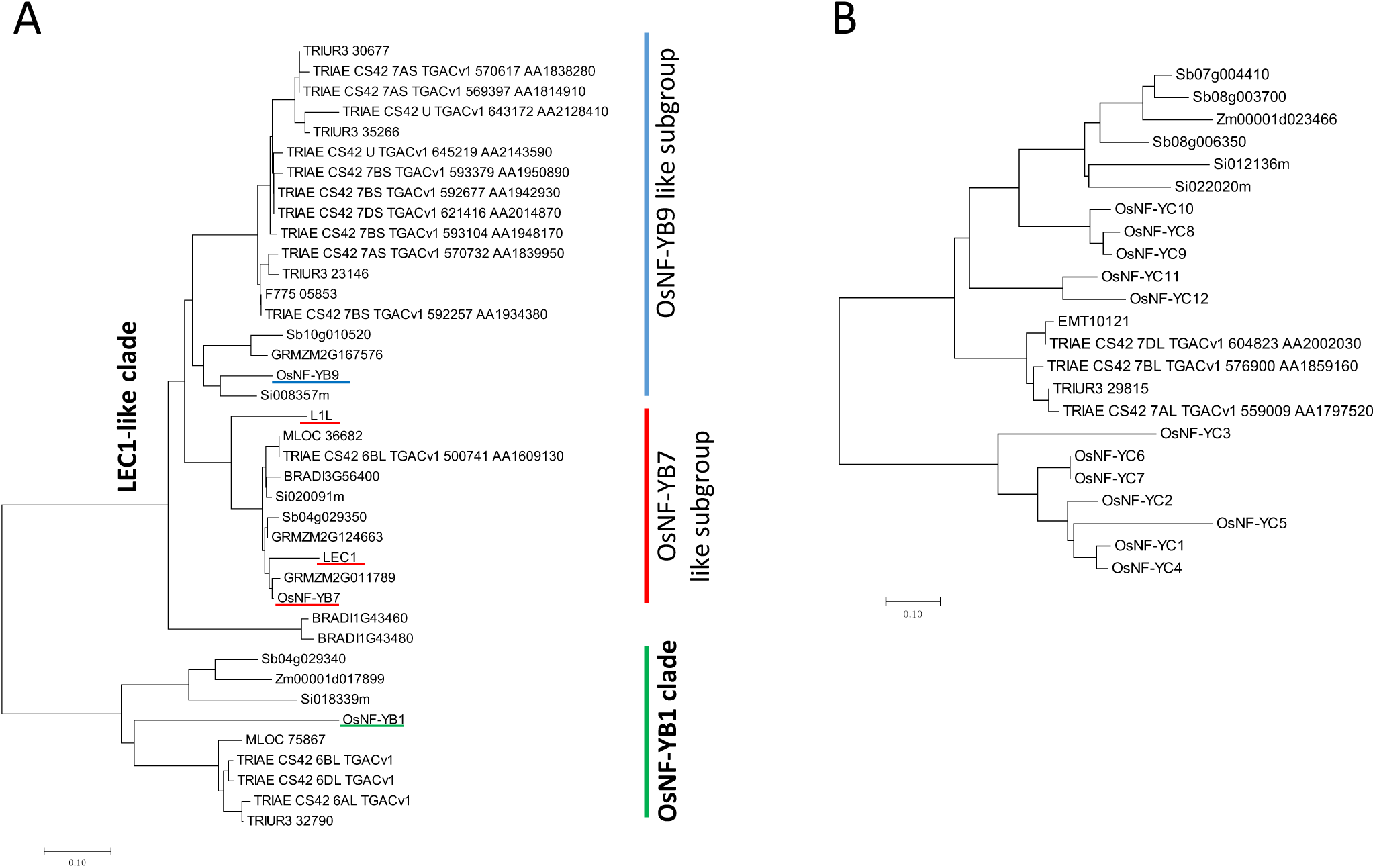
Neighbor-joining phylogenetic trees of the endosperm-preferential OsNF-YBs and OsNF-YC. **(A)** Phylogenetic tree of the OsNF-YB1, OsNF-YB7, OsNF-YB9 and their homologs. **(B)** Phylogenetic tree of the OsNF-YCs and the close homologs of OsNF-YC8 and OsNF-YC12. The locus identities initiating with TRIUR, TRIAE, F775, Sb, GRMZM(ZM), Os, Si, At, MLOC and BRADI indicate that genes are from *Triticum urartu, Triticum aestivum, Aegilops tauschii, Sorghum bicolor, Zea mays, Oryza sativa, Setaria italica, Arabidopsis thaliana, Hordeum vulgare* and *Brachypodium distachyon*, respectively. The core sequences of the conserved domains of NF-YBs and NF-YCs were used for trees construction.

### Endosperm-preferential expression of NF-Ys in other monocot species

Since the rice endosperm-preferential NF-Ys had variability in their core sequences, we further determined if such variability is present in NF-Y members of other plant species. Zm00001d017899 and Sb04g029340 are the most similar homologs of OsNF-YB1 in maize and sorghum (**Fig. 3A**), respectively. By searching RNA-seq data deposited in the Expression Atlas database, we found that these genes are predominantly or exclusively expressed in the endosperm (**Fig. 4A**). Likewise, Zm00001d045772 (GRMZM2G167576) and Sb10g010520, the maize and sorghum homologs of *OsNF-YB9* (**Fig. 3A**), are also exclusively expressed in the endosperm (**Fig. 4B**). Sb08g006350, Sb08g003700, and Sb07g004410 are three homologs of the endosperm-preferentially expressed rice NF-YCs in sorghum (**Fig. 3B**), and we found that the mRNA transcripts of Sb08g006350 and Sb07g004410 were only detectable in the endosperm (**Fig. 4B**). The expression profile of Sb08g003700 is not available in the Expression Atlas. However, with the data availability in MOROKOSHI Sorghum Transcriptome Database, it was observed that Sb08g003700 specifically expressed in the endosperm **(Supplementary Fig. 4A).** The maize genome encodes only one homology gene, Zm00001d023466 (GRMZM2G052499), of *OsNF-YC8* or *OsNF-YC12.* Again, the Expression Atlas had no information about this gene, but the microarray-based experiments indicated that Zm00001d023466 was predominantly expressed in the endosperm **(Supplementary Fig. 4B).** Taken together, these results clearly indicate that monocots have evolved a group of phylogenically conserved NF-YBs and NF-YCs that are predominantly expressed in the endosperm. This may occur after the divergence of dicots and these genes most likely involved in the regulation of endosperm development. We also explored the expression pattern of *OsNF-YB7* homologs in other monocot species. Similar to the *OsNF-YB7*, Zm00001d017898 (GRMZM2G011789) and Zm00001d051697 (GRMZM2G124663) of maize and Sb04g029350 of sorghum, were predominantly expressed in the embryo (**Fig. 4, Supplementary Figs. 4C and D),** indicating that these genes may function in seed development similar as the role of *LEC1* in *Arabidopsis.*

**Figure 4.**
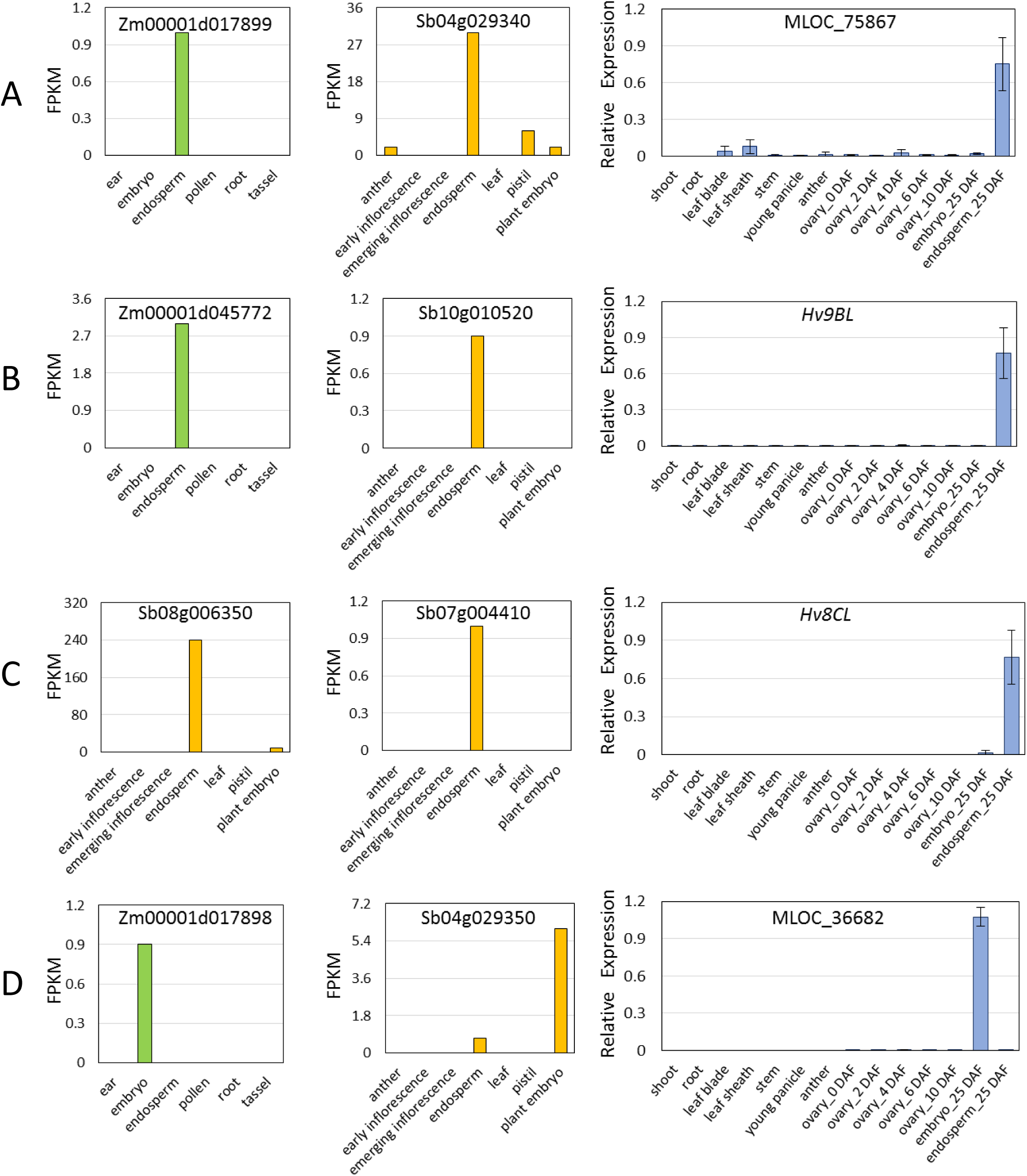
Gene expression of the *OsNF-YB1 Likes* (A), *OsNF-YB7 Likes* (B), *OsNF-YB9 Likes* (C) and *OsNF-YC8/12* Likes (D) in maize, sorghum and barley. The green, yellow and blue bars indicate gene expression of the maize, sorghum and barley homologs, respectively. The expression abundance in maize and sorghum are indicated by FPKM (fragments per kilobase of exon per million fragments mapped). Data is obtained from the Expression Atlas. The relative expression in barley is detected by Real-time PCR assay. Y-axes indicate the tissues used for expression analysis. DAF, days after fertilization. Three biological replicates were used for analysis; error bars indicate standard deviations.

The homologs of OsNF-YB1 and OsNF-YB7 were found in barley (**Fig. 3A)** through protein balst against the protein dataset of barley. However, we did not find the homologs of OsNF-YB9, OsNF-YC8 or OsNF-YC12, possibly due to poor annotation of the barley genome. Instead, when we used the core protein sequences of these rice NF-Ys to search against the barley genomic DNA sequence, two genes were discovered (referred as *Hv9BL* and *Hv8CL* hereafter) that showed high similarities with OsNF-YB9 and OsNF-YC8/12 at the core sequence of the conserved domains **(Supplementary Fig. 5).** As expected, the MLOC_75867 (an *OsNF-YB1* homolog), *Hv9BL* and *Hv8CL* were predominantly expressed in the endosperm, whereas the MLOC_36682 (an *OsNF-YB7* homolog) showed high expression in the embryo but not in the endosperm (**Fig. 4**).

Sequence similarity and conserved expression pattern of these endosperm-preferential NF-Y groups in different monocot species strongly suggest that the functional differentiation of these genes occurred in a common rice, maize, and sorghum ancestor. Notably, *OsNF-YB1* (LOC_Os02g49410) and *OsNF-YB7* (LOC_Os02g49370) and their corresponding homology genes in maize (Zm00001d017899 and Zm00001d017898) and sorghum (Sb04g029340 and Sb04g029350), are located on adjacent chromosomal regions **(Supplementary Fig. 6).** The observation further confirms that the gene duplication event may responsible for the evolution of NF-YB1 and NF-YB7 is very ancient.

### Subcellular localization of the endosperm-preferentially expressed OsNF-Ys

To detect the subcellular localization of the endosperm-preferential NF-Ys, we N-terminally fused the NF-Ys with a Venus-tag and transiently expressed in tobacco (*Nicotiana benthamiana*) epidermal cells. The results indicated that NF-YA8 was predominately targeted to the nucleus (**Fig. 5A**), while the other endosperm-preferential OsNF-Ys were localized in, both, the cytoplasm and the nucleus (**Figs. 5B-H**). The results are consistent with previous findings that NF-YB and NF-YC can be dimerized in the cytoplasm and then be imported into the nucleus to form a heterotrimer with NF-YA (Laloum *et al.*, 2013).

**Figure 5.**
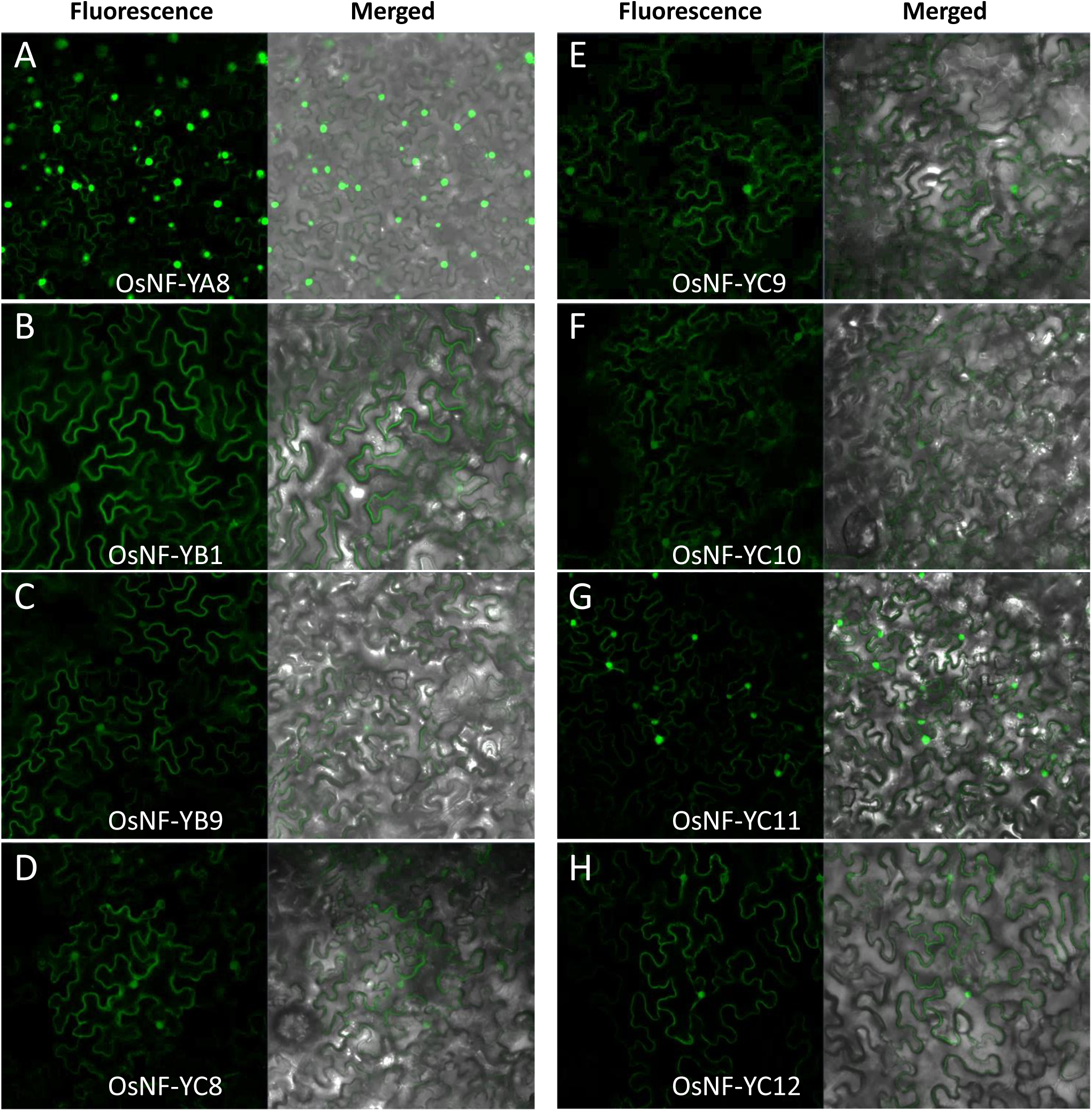
Subcellular localization of the endosperm-preferential NF-Ys of rice in tobacco epidermal cells. The florescence signal of the OsNF-YA8:Venus fusion is predominantly expressed in nucleus **(A)** while these of the rest OsNF-Y:Venus fusions are expressed in cytoplasm, as well as in nucleus **(B-H).** The NF-Ys are N-terminally fused to a venus tag.

### Interactions between the endosperm-preferential OsNF-Ys

The core sequences of endosperm-preferential NF-Ys are somewhat different from the canonical NF-Ys **(Supplementary Fig. 1);** however, the homology modeling analysis showed no change of the protein structures in spite of sequence variabilities **(Supplementary Fig. 7).** The results indicated that these endosperm-preferential NF-Ys may interact with different members as the canonical NF-Ys. To test this idea, we performed a yeast two-hybrid assay to detect protein interactions between different groups of endosperm-preferential NF-Ys of rice. The results showed that OsNF-YA8 was only able to interact with OsNF-YB9 (**Fig. 6A and Supplementary Fig. 8**). Notably, despite the high similarity between OsNF-YB7 and OsNF-YB9, OsNF-YA8 could not interact with OsNF-YB7 in yeast **(Supplementary Fig. 8).** On the other hand, all the NF-YBs tested, including OsNF-YB1, OsNF-YB9, and OsNF-YB7, were able to interact with all the endosperm-preferential NF-YC family members **(Supplementary Fig. 9).**

**Figure 6.**
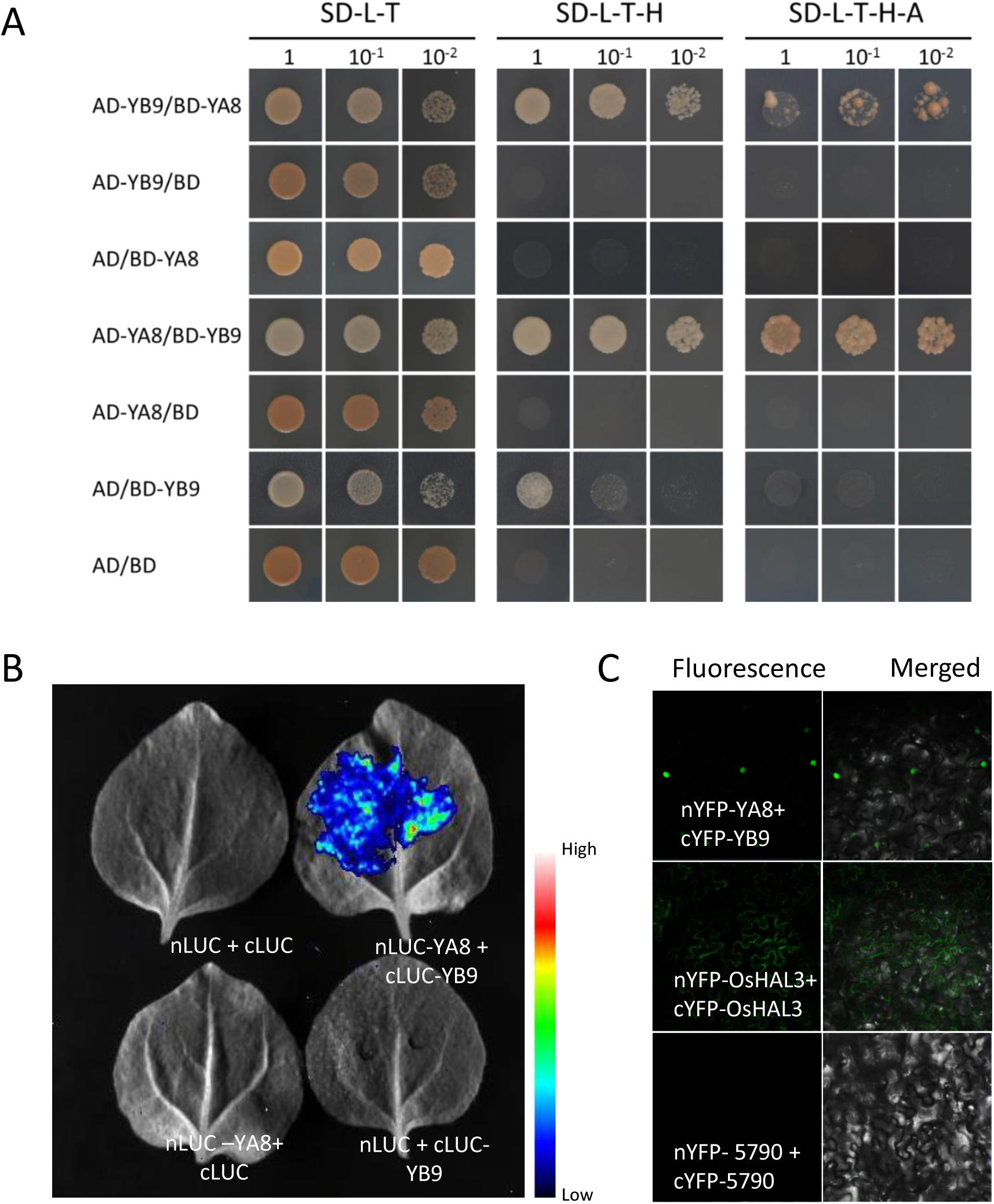
Interactions between the OsNF-YA8 and OsNF-YB9. **(A)** Yeast-two-hybrid assay shows interaction between OsNF-YA8 and OsNF-YB9. AD and BD indicate the activation domain and the DNA binding domain of GAL4, respectively; AD-YA8, AD-YB9, BD-YA8 and BD-YB9 indicate AD:OsNF-YA8, AD:OsNF-YB9, BD:OsNF-YA8 and BD:OsNF-YB9 fusions, respectively. Serial dilutions of the yeast cells expressing the indicated proteins were plated on the non-selective medium (SD-L-T) or selective plates (SD-L-T-H and SD-L-T-H-A). **(B)** Split luciferase complementation assay confirms the interaction between OsNF-YA8 and OsNF-YB9. nLUC, cLUC, nLUC-YA8 and cLUC-YB9 indicate the N-terminal of luciferase, the C-terminal of luciferase, the nLUC:OsNF-YA8 fusion and the cLUC:OsNF-YB9 fusion, respectively. Different constructs combinations are transiently co-expressed in the tobacco epidermal cells. **(C)** BiFC assay shows interactions between OsNF-YA8 and OsNF-YB9. OsNF-YA8 and OsNF-YB9 are fused with the N-terminal of YFP (nYFP) and the C-terminal of YFP (cYFP), respectively. OsHAL3, which can dimerize in the cytoplasm, is used as the positive control, while the gene LOC_Os01g05790 (5790), which cannot form a homo-dimer, is used as the negative control.

Usually, NF-YA does not interact with NF-YB or NF-YC monomer (Hackenberg et al., 2012). To confirm the interaction between OsNF-YA8 and OsNF-YB9, we conducted a split luciferase complementation assay in the tobacco epidermal cells. After the co-filtration of nLUC (the N-terminal of luciferase) and cLUC (the C-terminal of luciferase), we failed to detect luciferase activity (**Fig. 6B**). However, co-expression of nLCU-OsNF-YA8 and cLUC-OsNF-YB9 showed a high luciferase activity (**Fig. 6B**). In addition, neither the combination of nLCU-OsNF-YA8 and cLUC, nor the combination of nLCU and cLUC-OsNF-YB9 could activate luciferase activity (**Fig. 6B**). The results confirmed the interaction between OsNF-YA8 and OsNF-YB9 in plant. Next, we used a bimolecular fluorescence complementation (BiFC) assay to determine the organelle in which the interaction occurs. The fluorescence signal was observed in the nucleus but not in the cytoplasm (**Fig. 6C**), which is consistent with previous findings that NF-YA is a nuclear localized protein and can bind to a NF-YB/NF-YC dimer to form heterotrimer (Laloum *et al.*, 2013).

### Endosperm preferential NF-YBs can interact with ERFs

A previous study has indicated that OsNF-YB1 can interact with the ERF protein, OsERF115, to form NF-YB/NF-YC/ERF heterotrimers in the nucleus of rice endosperm (Xu *et al.*, 2016). We performed yeast-two-hybrid assays to test whether the other NF-YBs have the same ability. The results suggested that OsNF-YB9 was able to interact with OsERF114 and OsERF115 (**Fig. 7A)** but not with OsERF74 and OsERF72 **(Supplementary Fig. 10).** Notably, OsNF-YB7 also showed an interaction with OsERF115, and possibly with OsERF114 as well (**Fig. 7A**). We also tested the interaction between OsERF114/115 and the endosperm-preferential OsNF-YA8 and OsNF-YCs. None of the ERFs interacted with OsNF-YA8 (**Fig. 7B**). However, OsNF-YC12 could interact with OsERF114 in yeast (**Fig. 7B**). Due to the self-activation activity of OsNF-YC8 (**Fig. 7B**), the interactions between OsERFs and OsNF-YC8 need to be further confirmed. Interestingly, the expression patterns of *OsERF114* and *OsERF115* were very similar to those of the endosperm-preferential NF-Ys **(Supplementary Fig. 11),** indicating that OsERF114 and OsERF115 may coordinate with endosperm-preferential NF-YB and NF-YC members to regulate rice endosperm development.

**Figure 7.**
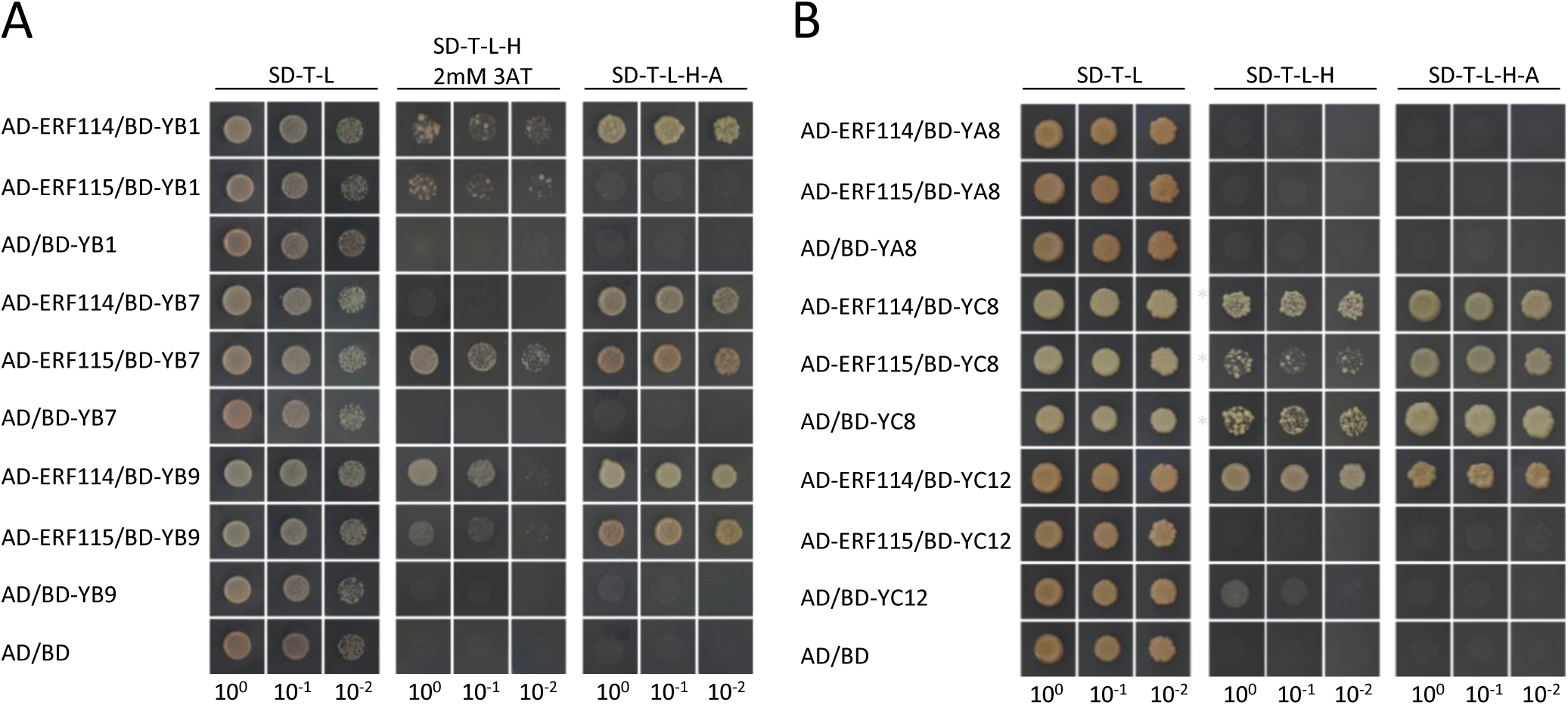
Interactions between the seed-preferential OsNF-Ys and ERFs. (A) OsNF-YB1, OsNF-YB7 and OsNF-YB9 interact with OsERF114 or OsERF115. BD-YB1, BD-YB9 and BD-YB7 indicate that the OsNF-YB1, OsNF-YB9 and OsNF-YB7 are C-terminally in fusion with the DNA binding domain of GAL4 (BD), respectively. AD-ERF114 and AD-ERF115 indicate that the OsERF114 and OsERF115 are C-terminally in fused with the activation domain of GAL4 (AD), respectively. Serial dilutions of the yeast cells expressing the indicated proteins were plated on the non-selective medium (SD-L-T) or selective plates (SD-L-T-H with 2mM 3AT or SD-L-T-H-A). (B) OsNF-YC12 interacts with OsERF114 in yeast. BD-YA8, BD-YC8 and BD-YC12 indicate that the OsNF-YA8, OsNF-YC8 and OsNF-YC12 are C-terminally in fusion with the BD, respectively. The yeast grew on the selective triple dropout medium with 5mM 3AT are indicated by stars. High self-activation activity of OsNF-YC8 was observed; the yeast cells expressing the AD and the BD:OsNF-YC12 fusion was survivable on the selective medium.

### Transcriptional activation activity

It is yet to determine whether the members of NF-Y family have any transcriptional activation activity in plants. We fused the endosperm-preferential NF-Ys of rice with the GAL4 binding domain to detect their transcriptional activation ability in yeast. OsNF-YA8, OsNF-YB1, OsNF-YB9, and OsNF-Y7B did not survive on the selective synthetic dropout medium **(Supplementary Fig. 12),** which suggested that these proteins alone are not capable of transcription activation. In terms of the NF-YC family members, OsNF-YC8, OsNF-YC9, and OsNF-YC10 could survive on the dropout medium like the positive control (the fusion of the GAL4 activation domain with the GAL4 DNA binding domain), whereas both OsNF-YC11 and OsNF-YC12 had not shown any transcriptional activation activity **(Supplementary Fig. 12).** Next, we fused the N-terminal (containing the core domain of NF-YC) of OsNF-YC8 with the C-terminal of OsNF-YC12 (designed as YC8N+YC12C hereafter), and found that this chimeric protein did not show transcriptional activity (**Fig. 8A**). In contrast, transformants with the fusion of the N-terminal of OsNF-YC12 and the C-terminal of OsNF-YC8 (YC12N+YC8C) could survive on the selective medium (**Fig. 8A**), which indicated that the C-terminal of OsNF-YC8 determines the transcriptional activation activity.

**Figure 8.**
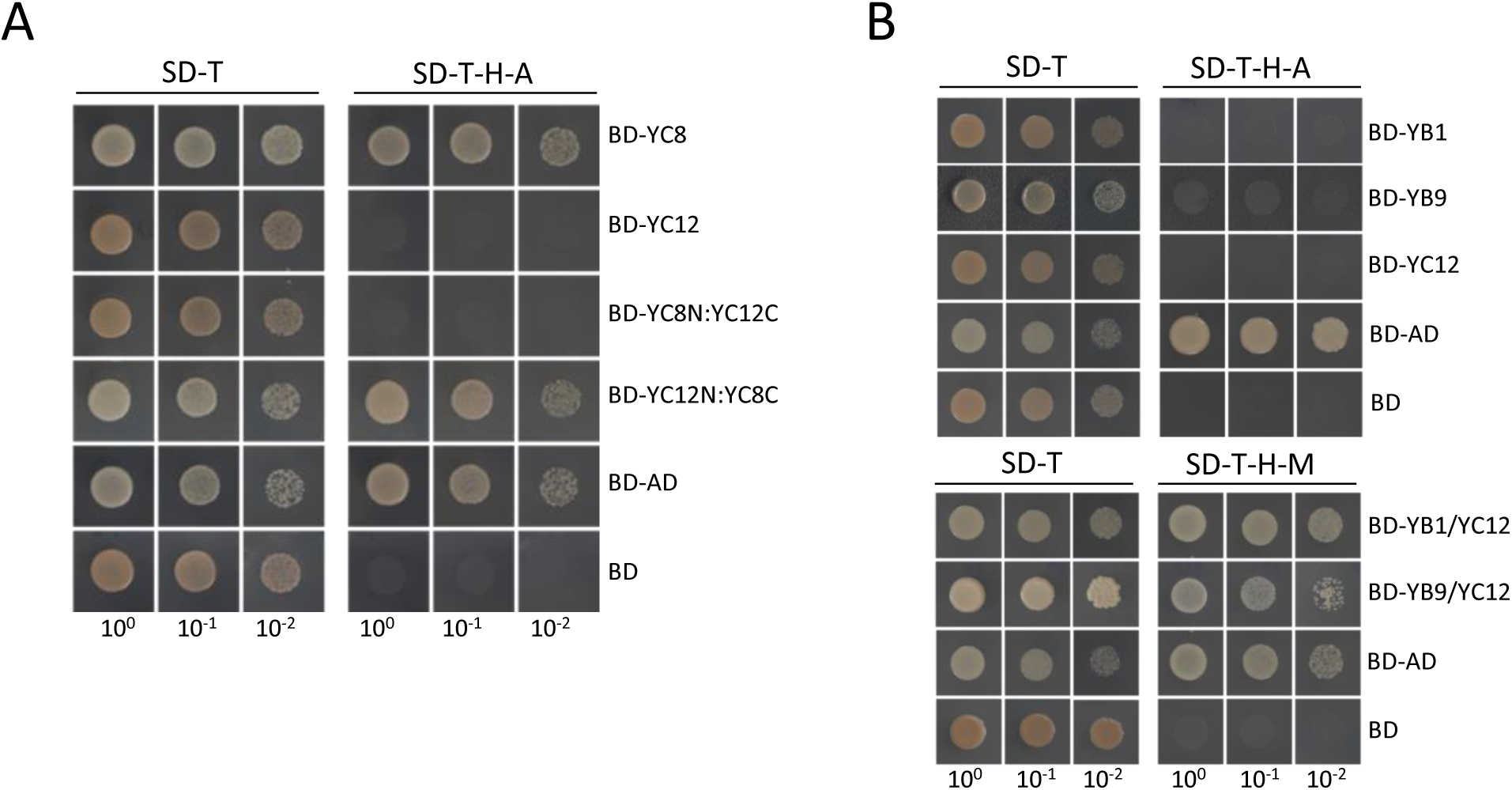
Transcriptional activation activities of the endosperm-preferential OsNF-Ys. **(A)** C-terminal of OsNF-YC8 is important for the transcriptional activation activity. BD and AD indicate the DNA binding domain and activation domain of GAL4, respectively. The serially diluted yeast cells expressing BD:OsNF-YC8 fusions (BD-YC8) and BD:OsNF-YC12 fusions (BD-YC12) could grow on the nonselective medium (SD-T), but only the transformants expressing BD-YC8 survived on the selective medium (SD-T-H-A), like what the BD:AD (a positive control) expressing yeast cells performed. When using the C-terminal of the OsNF-YC12 (amino acid residuals 361-984) to replace the C-terminal of OsNF-YC8 (385-1392), the fusions (BD-YC8N:YC12C) lose transcriptional activation activity. In contrast, the OsNF-YC12 N-terminal (1-360) and OsNF-YC8 C-terminal (385-1392) fusion (BD-YC12N:YC8C) showed transcriptional activation activity. **(B)** The OsNF-YB1 and OsNF-YC12 dimer shows transcriptional activation activity in yeast. The BD:OsNF-YB1, BD:OsNF-YB9 and BD:OsNF-YC12 do not show transcriptional activation activity; but the yeast cells simultaneously expressing of BD:OsNF-YB1 and OsNF-YC12 (BD-YB1/YC12), and BD:OsNF-YB9 and OsNF-YC12 (BD-YB9/YC12) could survive on the selective medium (SD-H-T-M). The OsNF-Ys in the upper and lower panel of **(B)** were cloned into pGBKT7 and pBridge, respectively.

Because OsNF-YC11 and OsNF-YC12 alone lacks transcriptional activation activity, we assumed that the transcriptional activation ability of these NF-YC family members may require the assistance of the NF-YBs it interacted with. OsNF-YB1 and OsNF-YC12 can interact with each other **(Supplemental Fig. 9A),** but neither of OsNF-YB1 nor OsNF-YC12 showed transcriptional activation activity (**Fig. 8B and Supplementary Fig. 12**). However, when we expressed the fusion of OsNF-YB1 and the GAL4 DNA binding domain, and OsNF-YC12 simultaneously in yeast, the transformants could survive on the selective dropout medium (**Fig. 8B**). The observation strongly supported our hypothesis that the formed dimer OsNF-YC12 and OsNF-YB1 exhibits transcriptional activation activity.

### Phenotypic analysis of the OsNF-Ys knockout mutants

To uncover the biological roles of the endosperm-preferential OsNF-Ys in seed development, the knockout mutants of these genes were generated. We took the advantage of CRISPR/Cas9 constructs that expressed the guide RNAs to target different regions of the first-half of the endosperm-preferential *OsNF-Ys.* Out of the 8 genes, we have obtained the homozygous T_0_ mutants of *OsNF-YB1, OsNF-YC8, OsNF-YC11* and *OsNF-YC12* to date **(Supplementary Fig. 13).** The mutants did not show any visible defects of vegetative development, which in agreement with their endosperm-preferential expression pattern. Consistent to previous findings (Sun *et al.*, 2014; Bai *et al.*, 2015; Xu *et al.*, 2016), the *osnf-yb1* mutant showed reduced seed size and increased chalkiness of endosperm (**Figs. 9A-C**). However, *osnf-yc8, osnf-yc11* and *osnf-yc12* did not display any seed abnormalities, in terms of the seed size and endosperm appearance (**Figs. 9A-C**). Most possibly, it is due to the redundancies among *OsNF-YC8, OsNF-YC9* and *OsNF-YC10* and between *OsNF-YC11* and *OsNF-YC12.* Notably, disruption of *OsNF-YB7* seemed not affect the endosperm development of the mutant as well (**Figs. 9A-C**).

**Figure 9.**
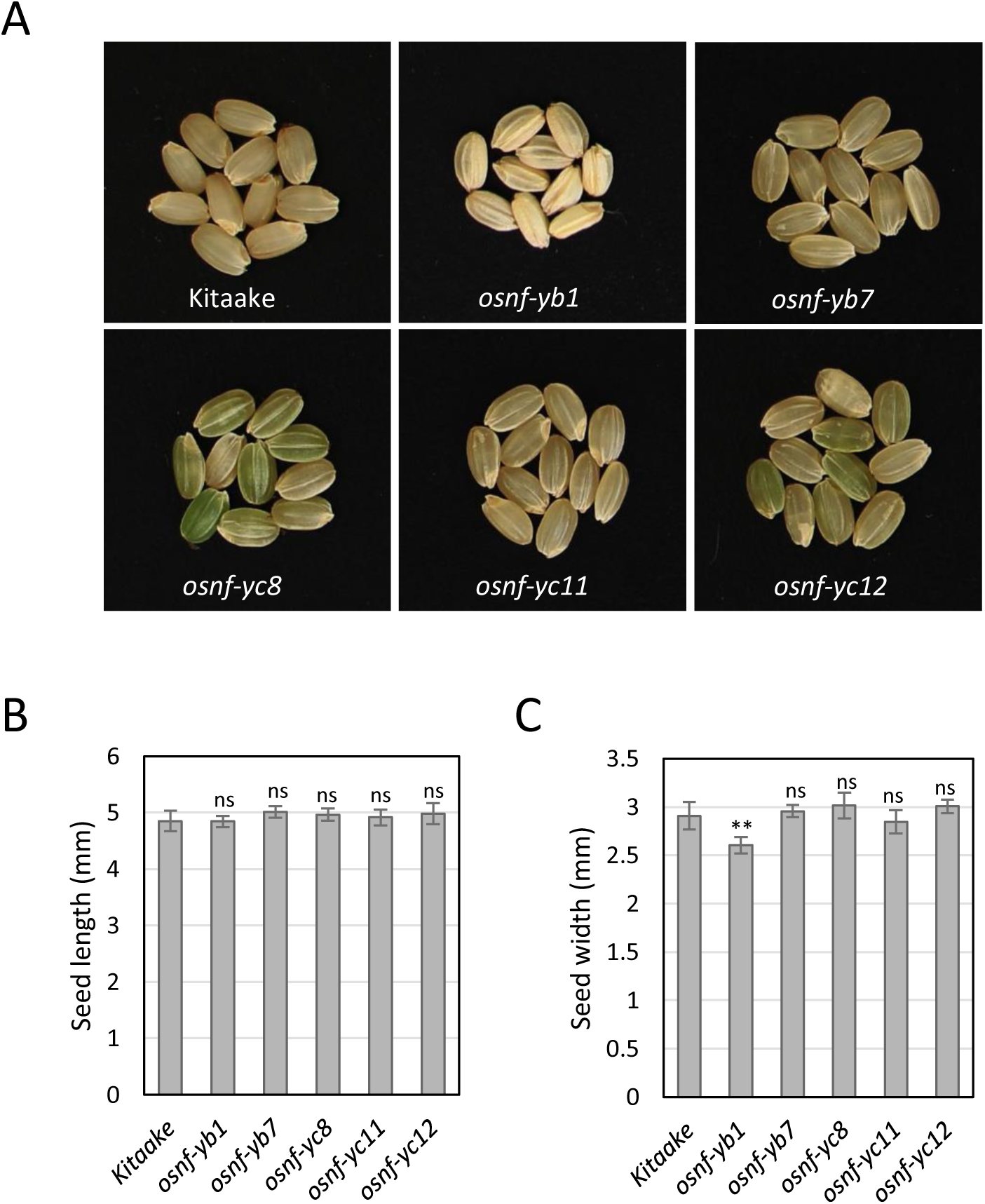
Seed morphologies of the seed-preferential *OsNF-Ys* mutants generated by CRISPR/Cas9 approach. **(A)** Seed morphology of the *osnf-yb1, osnf-yb7, osnf-yc8, osnf-yc11, osnf-yc12* and the wild type (Kitaake). **(B-C)** Seed length **(B)** and seed width **(C)** of the *osnf-yb1, osnf-yb7, osnf-yc8, osnf-yc11, osnf-yc12* and the wild type (Kitaake). More than 30 well-filled seeds from the main stem were chose for the measurements; error bars indicate standard deviations. **, *p*<0.01; ns, no significance; t-test was used for the statistical analysis.

## Discussion

Functional analyses revealed that plant NF-Ys may have multiple functions that regulate stress response, flowering time, embryo development, and chloroplast biogenesis (see review Petroni et al., 2012; Laloum et al., 2013). As compared to *Arabidopsis*, the function of the rice NF-Ys is much less understood. Although the expression of some NF-Y genes in the endosperm has been reported (Stephenson et *al.*, 2007; Yang *et al.*, 2016; Zhang *et al.*, 2016*a*), the biological function of plant NF-Ys on endosperm development is largely unknown. To explore the functions of NF-Ys in endosperm development, we conducted a comprehensive phylogenetic analysis of endosperm-preferential NF-Ys in rice. Our results suggest that there are several phylogenetically-conserved groups of NF-YB and NF-YC that may play critical roles in cereal endosperm development (**Fig. 3**). Moreover, these genes in other monocot species were also predominantly expressed in the endosperm (**Fig. 4**), which implies that the functional role of the group of genes in endosperms is much conserved in monocots. We could not find any homologs of these endosperm-preferential NF-Ys in dicots, indicating that the genes are evolved after the divergence of dicots and monocots. The fate of the endosperm in dicots and monocots is distinct (Zhou *et al.*, 2013). Dicots usually consume the endosperm for embryo growth, whereas monocots preserve the endosperm as an organ for starch and nutrient accumulation. We believe that the endosperm-preferential NF-Ys are most likely the candidates that may contribute in the divergence of monocots and dicots. Previous studies have shown that LEC1-family NF-YBs are essential for seed maturation and embryo development (Kwong *et al.*, 2003; Lee *et al.*, 2003). Interestingly, here we found that the LEC1-like proteins could be divided into two plant subgroups (**Fig. 3A**). The OsNF-YB7 and its homologs in other monocot species were phylogenically close to LEC1 (**Fig. 3A**). Resembling the expression pattern of *LEC1*, these genes were also predominantly expressed in the embryo (**Fig. 4 and Supplementary Figs. 4C and D).** OsNF-YB9 and its homologs in other monocot species consist of the other clade of LEC1-like proteins (**Fig. 3A**). Distinct from the *OsYB7Ls*, the *OsYB9Ls* of rice, maize, barley (*Hordeum vulgare*), and sorghum were endosperm-preferentially expressed (**Fig. 4 and Supplementary Fig. 4).** The findings suggest that some LEC1-like NF-YBs of monocots are sub-functionalized for endosperm development.

A systematic interaction analysis of different seed-preferential NF-Ys showed that OsNF-YA8 could interact with OsNF-YB9 but not with the tested members of NF-YB and NF-YC gene family (**Fig. 6 and Supplementary Fig. 8).** This was surprising, because NF-YAs usually binds to NF-YB/NF-YC dimers and not to any single subunit (Hackenberg *et al.*, 2012). The functional significance of this interaction in rice seed development is unclear and requires further studies. Notably, in spite of having high similarities at the core domain sequences, OsNF-YA8 failed to show any interaction with OsNF-YB7 in yeast **(Supplementary Fig. 8).** This raises the possibility of other sequences outside the core domain which might play a role in the interaction of OsNF-YA8 and OsNF-YB9.

In *Arabidopsis*, NF-YA family members showed variability in CCAAT-binding activity (Calvenzani *et al.*, 2012). A phylogenic analysis suggested that OsNF-YA8 was a recently-evolved gene that might be present only in the *Oryza* genus. Whether or not OsNF-YA8 binds to CCAAT-box is not known. OsNF-YA8 carries many variations within the core domain compared with the canonical NF-YA family members **(Supplementary Fig. 1B).** Several conserved residuals are differentiated in the A2 domain of OsNF-YA8, nevertheless the changes seemed not to alter protein structure **(Supplementary Fig. 7).** The A2 domain is involved in recognition and binding with the CCAAT motif (Gnesutta *et al.*, 2017). It remains to determine whether the sequence variations allow OsNF-YA8 to recognize *cis*-elements other than CCAAT-box. The CCAAT-binding ability of NF-YA requires NF-YB and NF-YC (Calvenzani *et al.*, 2012; Gnesutta *et al.*, 2017). In addition to NF-YAs, NF-YB and NF-YC dimers can interact with some other transcription factor family members to regulate downstream targets (Yamamoto *et al.*, 2009; Cao *et al.*, 2014; Huang *et al.*, 2015; Xu *et al.*, 2016). A previous study has showed that OsNF-YB1 was able to interact with OsERF115 to modulate grain filling in rice (Xu *et al.*, 2016). In the present study, we found that OsNF-YB9 and OsNF-YB7 of rice could also interact with OsERF114 or OsERF115 (**Fig. 7**). Moreover, The NF-YC family member OsNF-YC12 showed interaction with the OsERF114 as well. Similar to OsNF-Ys genes, the expression of *OsERF114* and *OsERF115* was observed endosperm specific which suggest that the interaction between ERFs and NF-Ys complex is an important event in regulating the endosperm development in rice **(Supplementary Fig. 11).** Therefore, we assumed that the interaction between ERFs and NF-Y complex is very important for the endosperm development of rice (**Fig. 10**).

**Figure 10.**
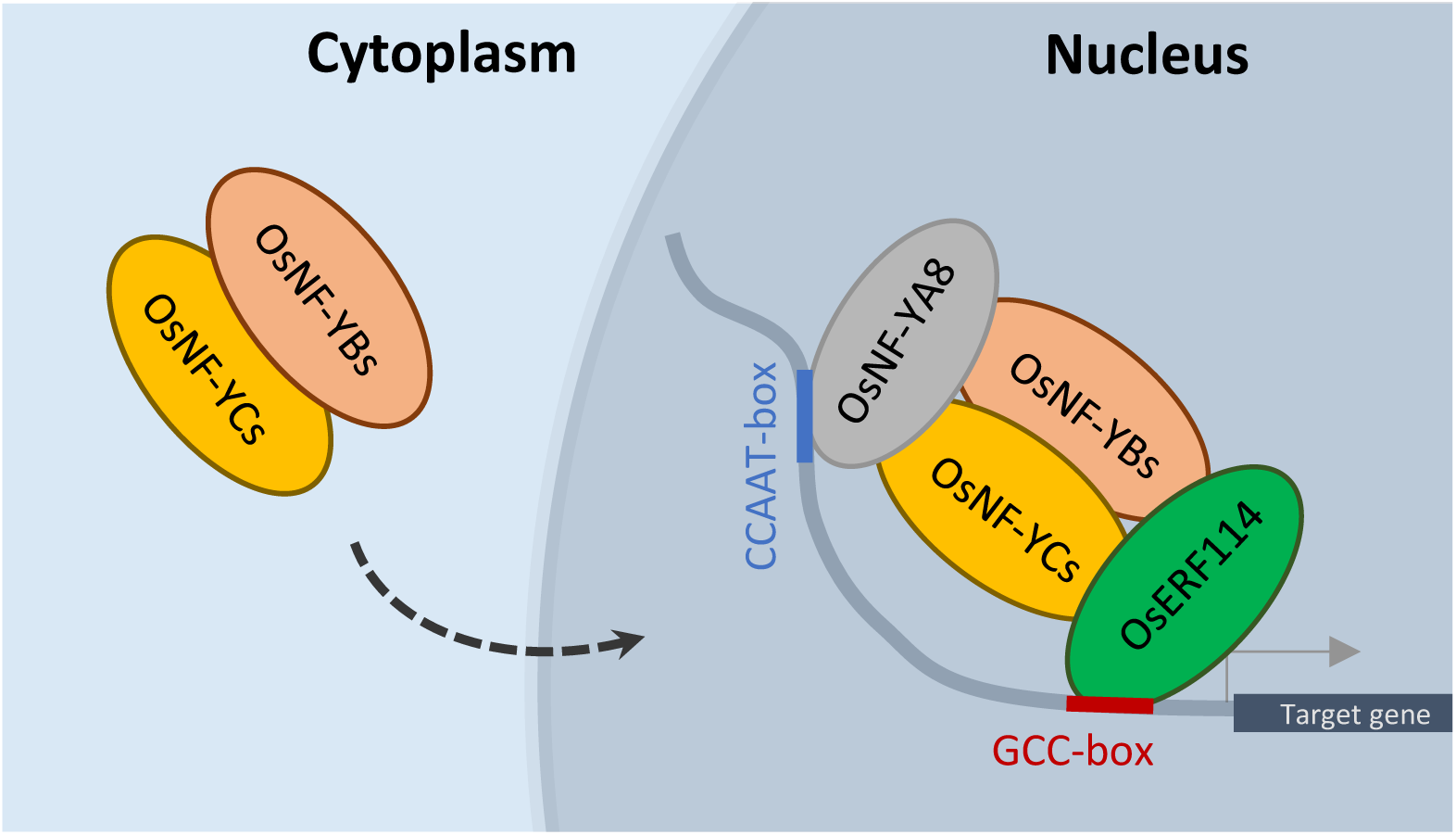
A hypothetical model showing the endosperm preferential OsNF-Ys function on rice endosperm development. The OsNF-YBs (OsNF-YB1/9 and possibly OsNF-YB7) and the OsNF-YCs (OsNF-YC8/9/10/11/12) are dimerized in the cytoplasm of endosperm cells and then imported into the nucleus to interact with OsNF-YA8 and OsERF114/115, or possibly with other transcription factors. OsNF-YA8 can recognize the CCAAT-motif while the OsERF114/115 can bind to the GCC-box. The OsERFs coordinate with the OsNF-Y complex to regulate the downstream genes involving endosperm development.

The previous mammalian studies have shown that the transcriptional activity of NF-Ys was dependent on the glutamine-rich domains which locate outside the core domains of NF-YB and NF-YC subunits (Coustry *et al.*, 1995, 1996). Our understanding of the transcriptional activation activity of plant NF-Ys is much poorer. Here, we found that, except OsNF-YC8 and its most similar rice homologs, OsNF-YC9 and OsNF-YC10, most of the endosperm-preferential NF-Y monomers in rice had no transcriptional activation activity in yeast **(Supplementary Fig. 12).** The C-terminal domain of OsNF-YC8 was essentially required for its transactivational activity (**Fig. 8A**). Interestingly, the dimer of OsNF-YB1/OsNF-YC12 displayed the transcriptional activation activity, though no such activity were observed in yeast by OsNF-YB1 or OsNF-YC12 alone (**Fig. 8B**). In addition, the OsNF-YB/OsNF-YC dimers can recruit other transcription factors, such as OsERF114, to form a more complicated complex to modulate the expression of downstream genes (**Fig. 10**). As an example, rice is likely to form a complex that consists of NF-Ys, Hd1, OsHAPL1, and the general transcription factors to control plant flowering (Zhu *et al.*, 2017). The NF-Y complex may also recruit transcription repressors to regulate gene expression (Li *et al.*, 2011). In this aspect, the NF-Y complex is more like a scaffold of transcriptional machinery. Leyva-González et al. (2012) showed that NF-YA2 of *Arabidopsis* could act like a repressor of a subset of genes that lack CCAAT-box. Therefore, the alternative interpretation of no transcriptional activation by OsNF-YA8, OsNF-YB1/7/9, or OsNF-YC11/12 alone in yeast could because these NF-Ys function as transcriptional repressors rather than activators. These findings suggested that the function of NF-Ys on transcriptional regulation might be very diverged, and the NF-Y complex of plants may require the coordination of multiple subunits for transcriptional activation activity. The role of endosperm-preferential NF-Ys in cereal endosperm development needs to be further discussed.

To further confirm the NF-Ys role in endosperm development, we generated knockout mutants of OsNF-Ys. Possibly due to functional redundancy among OsNF-Ys, we did not observe any seed phenotype with the knockout lines of *osnf-yc8, osnf-yc11* and *osnf-yc12* (**Figure 9**). However, the *osnf-yb1* displayed decreased seeds size and increased chalkiness of the endosperm (**Figure 9**). This is consistent with the previous findings (Sun *et al.*, 2014; Bai *et al.*, 2015; Xu *et al.*, 2016) and strongly implies that the OsNF-Ys play important functions in rice endosperm development. In summary, our findings indicated that the monocots have evolved a group of endosperm-preferentially expressed and phylogenetically-conserved NF-Ys in each family.

## Materials and Methods

### Plant materials and growth conditions

Kitaake (*O. sativa* subsp. *japonica*) rice plants were grown in a greenhouse with regular water and nutrient management. Various tissues, including the leaf blade, leaf sheath, flag leaf, stem, and young panicles at the booting stage were collected for RNA isolation. Before flowering, the plants were moved into a growth chamber that was maintained at a consistent 28 °C with 12 h/12 h light/dark cycle and 50% humidity. The seeds were labeled when flowering. Caryopsis of different ages were collected and immediately frozen by liquid nitrogen for RNA isolation. The barley variety Morex was planted in the experimental plot in 2016, Chengdu, China. Various barley tissues were collected at the flowering stage. The seeds were labeled on the day of flowering for sampling of different age caryopsis. Twenty-five days after fertilization, endosperms and embryos were carefully separated to avoid contamination of maternal tissues.

### Generation of the mutant plants

The targets for CRISPR/Cas9-mediated gene targeting were designed by the web-based tool CRISPR Primer Designer (http://plantsignal.cn/CRISPR/crispr_primer_designer.html). The targets were then cloned into the vector BGK032. All the constructs were transformed into the Agrobacterium strain EHA105 and used for plant transformation as described previously (Chen *et al.*, 2014). The DNA fragments embracing the targets of the plant transformants were amplified for sequencing. The T_0_ homozygous mutants of *osnf-yb1, osnf-yb7, osnf-yc8, osnf-yc11* and *osnf-yc12* were obtained for phenotypic analysis.

### RNA extraction and real-time PCR assay

Total RNA from different plant tissues was isolated using the Plant RNA Kit (OMEGA) following the manufacturer’s protocol. Total RNA was treated with an RNA-free DNase set (OMEGA). cDNA was synthesized by oligo-dT primers using PrimeScript RT Master Mix (TAKARA). The standard procedure provided by the manufacturer was used for the reactions. Two μL sample of the diluted cDNA was used for real-time PCR in a 20 μL reaction using the AceQ® qPCR SYBR® Green Master Mix (Vazyme, China). The real-time PCR reactions were performed on the CFX ConnectTM Real-Time System (BioRad). Three independent replicates were set for each sample. A rice proteasome gene (LOC_Os03g63430) was used as an internal control. Quantification of the relative expression was calculated by the ΔΔCt method. Primers used for the real-time PCR reactions can be found in **Supplementary Table 1.**

### Expression analysis of the endosperm-preferential NF-Ys in monocots

We used expression data deposited in Genevestigator® for the expression analysis of rice NF-Ys. A hierarchical clustering analysis of the expression of rice NF-Ys was also performed with the similarity search tool provided by Genevestigator®. Expression data of the *OsNF-YB1,7,9-like* and *OsNF-YC8,11-like* genes of maize and sorghum were obtained from the Expression Atlas (https://www.ebi.ac.uk/gxa/home). For those genes with no data in the Expression Atlas, we searched their expression through the Maize eFP Browser (http://bar.utoronto.ca/efp_maize/cgi-bin/efpWeb.cgi) for maize or the MOROKOSHI Sorghum transcriptome database (http://sorghum.riken.jp/morokoshi/Home.html) for sorghum.

### Alignment, phylogenetic analysis, and protein structure modeling

The core domains of OsNF-YA8, OsNF-YB1,7,9, and OsNF-YC8,11 were used as queries for Blast to identify their close homologs in other plant species. The protein and DNA sequences of the endosperm-preferential NF-Ys were obtained from the BioMart of the Phytozome database (www.phytozome.jgi.doe.gov/biomart/martview/). Alignment was conducted by Clustal Omega (www.ebi.ac.uk/Tools/msa/clustalo/). The phylogenetic neighbor-joining trees were generated by MEGA7.0 (Kumar *et al.*, 2016). The intensive model of the Phyre2 web portal was used for NF-Ys modeling (Kelley *et al.*, 2015).

### Yeast two-hybrid assay

The coding sequences of *OsNF-YA8, OsNF-YB1,7,9, OsNF-YC8,9,10,11,12*, and *OsERF72,74,114,115* were amplified by high-fertility DNA polymerase GXL (Takara) from endosperm cDNA using the primers list in **Supplementary Table 1.** The genes were cloned into either pGBK-T7 or pGAD-T7 using the ClonExpress® II One Step Cloning Kit (Vazyme). The prey and bait plasmids were transformed respectively into the yeast strains Y187 and Y2HGold. After mating of the two strains, co-transformants were selected on SD-L-T plates. Interactions were tested using SD-L-T-H with an optimized content of 3-AT and SD-L-T-H-A medium, simultaneously.

### Transcriptional activation assay in yeast

The coding sequences of *OsNF-YA8, OsNF-YB1,7,9*, and *OsNF-YC8,9,10,11,12* were cloned into pGBK-T7. Meanwhile, we amplified the GAL4 activation domain from a pGAD-T7 empty vector and fused it with GAL4 DNA-binding domain using the ClonExpress® II One Step Cloning Kit (Vazyme, China). The construct was used as a positive control for transcriptional activation activity. Meanwhile, we respectively cloned *OsNF-YB1* and *OsNF-YC12* into the multiple cloning site 1 (MCS1) and the MCS2 of the vector pBridge (Clontech). All of the constructs were then transformed into yeast strain Y2HGold to investigate the activity of transcriptional activation on the SD-Trp/-His/-Ade medium. The primers used in the experiment are listed in **Supplementary Table 1.**

### Sub-cellular localization analysis

The coding sequences of *OsNF-YA8, OsNF-YB1,7,9*, and *OsNF-YC8,9,10,11,12* were cloned into the binary vector p1300-LV using the ClonExpress® II One Step Cloning Kit (Vazyme). The constructs were transformed into *Agrobacterium tumefaciens* strain GV3101 and infiltrated into tobacco leaf epidermal cells to transiently express the NF-Y:Venus recombinant fusions. About 48 h after infiltration, the fluorescence signals were examined with a confocal laser scanning microscope (LSM710, Zeiss). Primer information is listed in **Supplementary Table 1.**

### BiFC and split luciferase complementation assay

The coding sequence of *OsNF-YA8* and *OsNF-YB9* was, respectively, cloned into pCAMBIA1300S-YN and pCAMBIA2300S-YC to fuse with the N-terminal and C-terminal of yellow fluorescence protein (nYFP and cYFP) and transformed into *Agrobacterium tumefaciens* strain GV3101. In addition, HAL3, which can dimerize in cytoplasm, was cloned into pCAMBIA1300S-YN and pCAMBIA2300S-YC as a positive control (Su *et al.*, 2016). The constructs, *nYFP-YA8* and *CYFP-YB9, nYFP-OsHAL3* and *cYFP-OsHAL3*, and *nYFP-5790* and *cYFP-5790* were co-infiltrated in tobacco leaves for about 48 h. The YFP fluorescence signals were examined with a confocal laser scanning microscope (LSM710, Zeiss).

Similarly, *OsNF-YA8* and *OsNF-YB9* was cloned respectively into binary vectors JW771 and JW772 to fuse with the N-terminal of luciferase (nLUC) and the C-terminal of luciferase (cLUC). *nLUC-OsNF-YA8* and *cLUC-OsNF-YB9* along with the combinations of *cLUC* and *nLUC, nLUC-OsNF-YA8* and *cLUC*, and *nLUC* and *cLUC-OsNF-YB9* were transiently expressed in tobacco leaves for about 48 h. The Luciferase Assay System (Promega) was used to detect the interactions. The chemiluminescence signal was detected by the Tanon Imaging System (5200 Multi, Tanon).

## Acknowledgements

This research was supported by grants from the National Key Research and Development Program of China (2016YFD0100902), the National Natural Science Foundation of China (31571623), the Natural Science Foundation of Jiangsu Province (BK20150446), and the Priority Academic Development of Jiangsu Higher Education Institutions.

## Supplementary materials

**Supplementary Figure 1.** Amino acid sequence alignment and phylogenetically analysis of the conserved domains of rice NF-Ys.

**Supplementary Figure 2.** Sequence logos of the conserved domains of OsNF-YB7 likes (OsYB7Ls) and OsYB9Ls.

**Supplementary Figure 3.** Multiple sequence alignment of the endosperm-preferential OsNF-YCs.

**Supplementary Figure 4.** Expression of some *OsNF-YC8 like* and *OsNF-YB7 like* genes in sorghum and maize.

**Supplementary Figure 5.** Multiple sequence alignments of the OsNF-YB9 and HvB9L (A) and of the OsNF-YC8, OsNF-YC12 and HvC8L (B).

**Supplementary Figure 6.** Diagram showing the physical linkage of the *OsNF-YB7 like* and the *OsNF-YB1 like* genes in rice, maize and sorghum.

**Supplementary Figure 7.** Predicted structures of the NF-Y conserved regions.

**Supplementary Figure 8.** OsNF-YA8 interacts with OsNF-YB9, but not with other endosperm-preferential OsNF-Ys.

**Supplementary Figure 9.** OsNF-YB1, OsNF-YB7 and OsNF-YB9 interact with all the endosperm-preferential OsNF-YCs.

**Supplementary Figure 10.** OsNF-YB1, OsNF-YB7 and OsYB9 do not show interaction with OsERF72 or OsERF74.

**Supplementary Figure 11.** *OsERF114* and *OsERF115* are activated after fertilization.

**Supplementary Figure 12.** OsNF-YC8, OsNF-YC9 and OsNF-YC10 exhibit transcriptional activation activities.

**Supplementary Figure 13.** The mutations in the mutants of endosperm-preferential OsNF-Ys.

**Supplementary Table 1.** The primers used in the present study.

